# Asymmetric phospholipids impart novel biophysical properties to lipid bilayers allowing environmental adaptation

**DOI:** 10.1101/2020.06.03.130450

**Authors:** Paul Smith, Dylan M. Owen, Christian D. Lorenz, Maria Makarova

## Abstract

Phospholipids are a diverse group of biomolecules consisting of a hydrophilic head group and two hydrophobic acyl tails. The nature of the head and length and saturation of the acyl tails are important for defining the biophysical properties of lipid bilayers. It has recently been shown that the membranes of certain yeast species contain high levels of unusual asymmetric phospholipids, consisting of one long and one medium chain acyl moiety – a configuration not common in mammalian cells or other well studied model yeast species. This raises the possibility that structurally asymmetric phospholipids impart novel biophysical properties to the yeast membranes. Here, we use atomistic molecular dynamics simulations (MD) and environmentally-sensitive fluorescent membrane probes to characterize key biophysical parameters of membranes formed from asymmetric lipids for the first time. Interestingly, we show that saturated, but asymmetric phospholipids maintain membrane lipid order across a wider range of temperatures and do not require acyl tail unsaturation or sterols to maintain their properties. This may allow cells to maintain membrane fluidity even in environments which lack the oxygen required for the synthesis of unsaturated lipids and sterols.

## Introduction

Membranes are composed of a richly diverse population of lipids. The primary class are glycerophospholipids which are themselves varied in their head group and the length and saturation of their two acyl tails. While cells can and do incorporate fatty acids from their environment into membrane phospholipids, they also synthesize and incorporate *de novo* fatty acids, expending energy and resources in the process and implying that compositional diversity is functionally relevant (Harayama and Riezman, 2018). The lipid composition of a bilayer profoundly influences the biophysical characteristics of that membrane, and this is particularly true when considering the length and saturation of the lipid acyl tails. The bilayer biophysical properties in turn influence membrane function – the diffusion and interactions of membrane proteins for example, or the curvature and bending rigidity during membrane remodeling.

One of the key biophysical properties influencing function is membrane lipid order, closely related to the phase behavior of the bilayer. In general, bilayers composed of glycerophospholipids can adopt one of two phases – a solid, or gel phase as low temperatures and a liquid-disordered phase at high temperatures (Heberle and Feigenson, 2011). Such a phase transition would be detrimental for a living organism living at variable temperatures and therefore cells incorporate sterols into the bilayer to modulate the phase behavior, allowing them to generate a new phase – the liquid-ordered phase, and maintain that phase over physiological temperature ranges. In general, the liquid-ordered phase is enriched in saturated phospholipids and sterols, whereas unsaturated phospholipids form lower order domains. The maintenance of membrane biophysical properties by cells using a combination of saturated lipids, unsaturated lipids and sterols, however, comes at a cost – unsaturated lipids and sterols require oxygen for their synthesis (Kwast et al., 1998; Shanklin and Cahoon, 1998).

In general, due to the characteristics of the lipid metabolic pathways responsible for their synthesis, the two acyl tails of each lipid are often similar or identical in length (Jenni et al., 2007; Lynen et al., 1980; Maier, 2006). For example, two of the most common lipids in mammalian cell membranes are di-oleoyl-phosphatidylcholine (DOPC) and di-palmitoyl-phosphatidylcholine (DPPC). It has recently been discovered that fission yeast species, *Schizosaccharomyces japonicus,* produces high quantities of structurally asymmetric phospholipids not common for mammals or even in closely related yeast species such as the well-known model organism *Schizosaccharomyces pombe* (Makarova et al., 2020). Curiously, *S. japonicus* shows several interesting environmental adaptations such as the ability to survive and proliferate at temperatures above 40 °C and in hypoxic environments (Kaino et al., 2018). Moreover, nuclear envelope remodeling during mitosis diverged between the two species with *S. japonicus* undergoing a semi-open mode and *S. pombe* executing closed mitosis (Makarova et al., 2016). This suggests that these membrane associated processes might be related to their distinct membrane composition and the resulting bilayer properties.

It is not currently known what biophysical properties membranes composed of asymmetric lipids will display but such an understanding is crucial to the future elucidation of their physiological role. Therefore, we investigate the biophysical properties of artificial lipid bilayers composed of these asymmetric phospholipids for the first time, using a combination of advanced fluorescence microscopy and atomistic scale molecular dynamics (MD) simulations. In particular, we form giant unilamellar vesicles from 1-stearoyl-2-decyl-sn-glycero-3-phosphatidylcholine (SDPC) which contains one saturated tail of 18 carbon length and one saturated tail of 10 carbon length. Vesicles are formed in the presence or absence of 30% ergosterol – the primary sterol found in fungi, and stained with the environmentally sensitive probe di-4-ANEPPDHQ which reports on membrane lipid order through changes in its fluorescence emission (Owen et al., 2012). Combined with insights from MD, our data shows that membranes formed from asymmetric lipids can maintain bilayer fluidity over a wide range of temperatures and, crucially, without the requirement for acyl tail double bonds or membrane sterols. We hypothesise that this might be a new and novel mechanism for the temperature and hypoxic tolerance of S*. japonicus* and we further propose bilayers composed of asymmetric lipids might have important medical and industrial applications due to the novel biophysical properties the asymmetric lipids impart.

## Materials and Methods

### GUV preparation

GUV preparation was performed by electroformation as previously described (Morales-Penningston et al., 2010). Briefly, a lipid film was formed on ITO-coated glass slides from 1 mg/ml of either a single lipid solution in chloroform or a phospholipid mixture with 30% (mol/mol) ergosterol. Asymmetric 1-stearoyl-2-decyl-sn-glycero-3-phosphatidylcholine (SDPC) was obtained through customized synthesis by Avanti Polar Lipids. After lipid films were dried, a 200 mM sucrose solution was used to form GUVs in the following conditions − 50°C, 11 Hz, 1V alternative electric current.

### Microscopy

Prepared GUVs were incubated with the 5 mM di-4-ANEPPDHQ and transferred to a glass-bottomed microscope dish. Imaging was performed in temperature-controlled conditions in the environmental chamber on a Zeiss LSM 780 inverted confocal microscope equipped with a 32 element GaAsP Quasar detector. The following parameters were used: a 488 nm laser was selected for fluorescence excitation of di-4-ANEPPDHQ and emission was detected for the ordered channel (500 – 601nm) and the disordered channel (640 – 700nm).

### Image analysis

GUV images obtained at different temperatures were analyzed using ImageJ and a custom written macro (Owen et al., 2012). Equatorial images of individual GUVs were selected for analysis and a mean GP value calculated for each GUV pixel. For each temperature point at least 20 individual GUVs of various radius (between 1 – 30μm) were analyzed.

### Molecular Dynamic Simulations

Six different lipid bilayers were simulated in order to investigate the effect of lipid molecules with asymmetric tails on the properties of the bilayers: pure DOPC, pure DSPC, pure SDPC, and mixed membranes containing each of these three lipids with 30% ergosterol. Each membrane was built with 200 total lipids (PC or PC & ergosterol) in each leaflet using the CHARMM-GUI membrane builder (Jo et al., 2008, 2009). Each system also included 30 water molecules per lipid molecule as well as 0.15 mM NaCl. Each membrane was minimised and then equilibrated to a temperature of 303.15 K and a pressure of 1 bar following the simulation protocol prescribed by CHARMM-GUI (Lee et al., 2016). After equilibrating each system, a production simulation was carried out for 200 ns at a temperature of 303.15K and a pressure of 1 bar. The temperature was controlled by a Nosé-Hoover thermostat and a Parrinello-Rahman barostat was used to control pressure.

All simulations were run using the GROMACS simulation package (van der Spoel and Hess, 2011) and the CHARMM36 forcefield was used to model the interactions of the lipid molecules and the ions (Klauda et al., 2010), while the water molecules were modelled using CHARMM TIP3P (Impey and Klein, 2016). The model for the SDPC lipid was generated by truncating the sn-2 tail of the lipid model used to represent DSPC after the 10^th^ carbon. LINCS constraints were used on the hydrogen-containing bonds in order to allow us to use 2.0 fs timesteps within the production simulations. Unless specified otherwise, the last 150 ns of production simulation was used for analysis.

The bilayer properties were characterised by the area per lipid (APL), lipid order parameter (S_CD_), bilayer thickness and the interdigitation of the lipids within the bilayers. Unless otherwise noted, analysis scripts were written in Python with the use of MDAnalysis (Gowers et al., 2016; Michaud-Agrawal et al., 2011).

A 2D Voronoi tessellation of atomic positions in each leaflet was performed to determine APL of each component of the bilayer, using the C21 C2 and C31 atoms as seeds for the PC lipids, O3 atoms for the ergosterol (atoms are given by CHARMM atom names). The lipid order parameter is a measure of the conformational flexibility of acyl chains in a bilayer, and is given by:

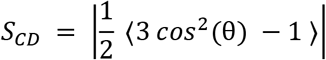

where θ is the angle between the bilayer normal and the carbon-hydrogen vector of a carbon atom in an acyl tail, and the average is taken over time and over all molecules of a given species within the membrane. The S_CD_ was calculated for each lipid or surfactant species a function of carbon atom position along an acyl chain. Smaller values of S_CD_ indicate a more disordered acyl chain. The bilayer thickness is calculated as distance between the mean height in z, along the bilayer normal, of the phosphate phosphorous atoms in the two leaflets.

In order to investigate whether the sn-1 hydrocarbon tails of the PC lipid molecules in the various bilayers ‘snorkel’ towards the water/bilayer interface, we measure the difference in the z-coordinates of the middle of the bilayer and the terminal carbon in each hydrocarbon tail of the lipid molecules. If the terminal carbon penetrates the opposite leaflet by more than 5Å then we count it as interdigitating with the other leaflet. If the terminal carbon is greater than 5Å away from the middle of the bilayer and still within the same leaflet as the headgroup of the lipid molecule then we count it as ‘snorkeling’. Otherwise we count it as being in the middle of the bilayer.

Lateral diffusion was calculated using the *gmx msd* module from the GROMACS package (Spoel and Hess, 2011) Using the Einstein relation, the diffusion coefficient (*D*) was evaluated from the mean square displacement (MSD):

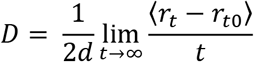

where *d* is the system dimension, *r* is the coordinate of the atom selection at a given time, *t*, from a time origin, *t*_*0*_.

## Results

### Membranes formed from asymmetric lipids display intermediate membrane lipid order consistent over physiological temperatures

We first generated GUVs composed of asymmetric phosphatidylcholine (SDPC) which were stained with 5μM di-4-ANEPPDHQ, imaged by confocal microscopy and the per pixel GP values calculated. For comparison, GUVs were also formed from pure symmetric saturated and unsaturated phospholipids. Figure 1A shows representative images of these GUVs imaged at physiological temperature (37°C) and pseudocoloured by GP value. Quantification of such GUVs per condition (Figure 1B) showed, as expected, that GUVs formed from symmetric unsaturated and saturated lipid show large negative and positive GP values, respectively. Interestingly, while SDPC is also a saturated lipid, it shows intermediate GP values, significantly different from both symmetric cases (p < 0.0005). This gives the first indication that asymmetric lipids may represent a mechanism to maintain some membrane fluidity in the absence of acyl chain double bonds.

**Figure 1:**
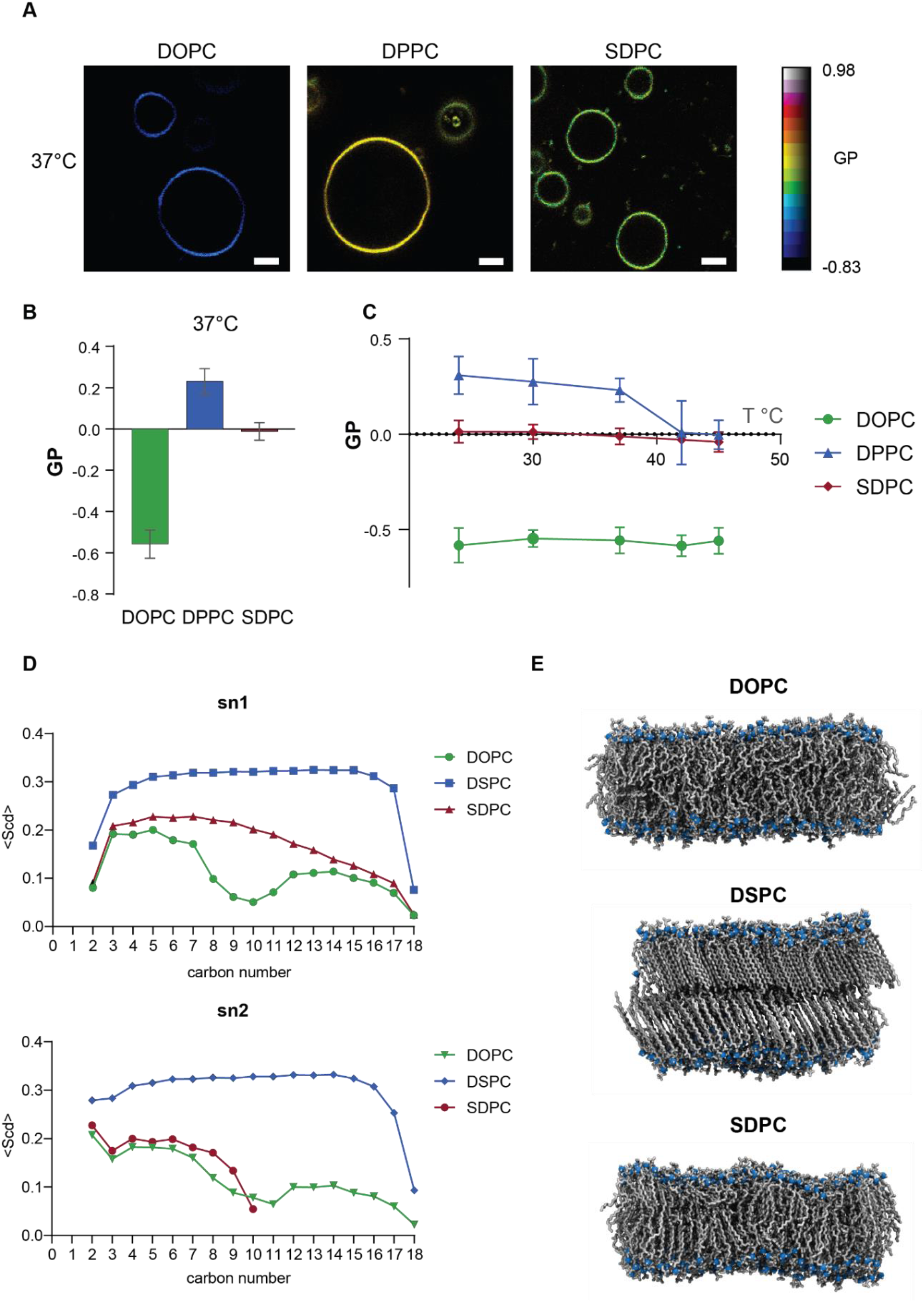
Lipid order of asymmetric phospholipid bilayers. A) Representative GP images of asymmetric and symmetric lipid GUVs stained with di-4-ANEPPDHQ. Color bar designates GP values (membrane lipid order). Scale bar, 5 μm. B) Quantification of membrane lipid order showing intermediate character of SDPC bilayers at 37°C. C) GP values over the range of 21°C −45°C. Note that the asymmetric lipids maintain membrane over a wide range of physiological temperatures. D) Deuterium NMR order parameter for the sn-1 (upper graph) and sn-2 (lower graph) chains. Note for sn-1 chain each bilayer shows intermediate character of SDPC and the presence of high order at the interfacial region and but low order in the bilayer core. E) Snapshots of atomistic MD simulations of the three bilayers.

We next examined the temperature dependence of these GP values. Both asymmetric SDPC and symmetric DOPC show relatively consistent levels of membrane order over the temperature range between 21°C and 45°C whereas the symmetric saturated lipid bilayers undergo a phase transition at around 42°C (Figure 1C). Interestingly GP values of asymmetric bilayers are similar to those of symmetric saturated lipids for temperatures above 42°C (Figure 1C).

To further detail the effect of asymmetry on membrane order, we performed atomistic MD simulations and calculated the deuterium NMR lipid order parameter. The order parameter for carbons on the sn-1 shows that the long tail of SDPC broadly behaved as a saturated lipid, without the central dip at the double bond position that DOPC displays (Figure 1D, upper graph). However, the simulations show that order is higher near the SDPC head, the area co-occupied by the short sn-2 chain and drops off to low levels deep in the bilayer. Therefore, the intermediate character of asymmetric lipids is the result of an order gradient from high order interfacial regions to a low order bilayer core. A similar trend is observed for the acyl tail at the sn-2 position (Figure 1D, lower graph). Intermediate character of SDPC lipid bilayer is visually represented in snapshots of these simulations (Figure 1E).

### Ergosterol has less effect on bilayers composed of asymmetric lipids and is not required to maintain bilayer fluidity

Sterols constitute approximately 30% of the mammalian and yeast membranes depending on the species, organelle distribution and growth conditions. Sterols modulate the biophysical properties of cell membranes maintaining membrane fluidity at physiological temperatures. We repeated the characterization of membrane lipid order using imaging and MD simulations for bilayers composed of asymmetric phospholipids this time in the presence of 30% ergosterol, the dominant sterol found in fungi. Representative GP images of these GUVs stained with di-4-ANEPPDHQ are shown in Figure 2A. The quantification (Figure 2B) shows, as expected ergosterol modulates membrane lipid order, generating relatively uniform GP values for all three bilayers. Interestingly, when compared to the bilayers lacking ergosterol, those formed from asymmetric lipids were not substantially affected by the presence of 30% ergosterol compared to their symmetric counterparts (Figure 2C). Again, the effect of asymmetric phospholipids on temperature dependence is profound, in particular when considering how membrane order changes when temperature is varied from 21°C (Figure 2D). Membranes composed of either saturated or unsaturated symmetric lipids show a marked decrease in membrane order over the temperature range 21°C to 45°C, in particular, likely undergoing a phase transition to the liquid-disordered phase at around 40°C. Bilayers composed of SDPC and ergosterol however are better able to maintain lipid order, eventually becoming the most ordered of the three conditions at 45°C. Quantification of the deuterium NMR lipid order parameter for each carbon atom of the sn-1 acyl tail (Figure 2E upper graph) shows the different effect of ergosterol on asymmetric compared to symmetric lipids. In the bilayer core, ergosterol has a minimal effect on SDPC lipid order, confirming the minimal changes in GP observed from the microscopy data. Again, a similar trend is observed for the acyl tail at the sn-2 position (Figure 2E, lower graph). Representative atomistic MD simulations of these bilayers are shown in Figure 2E.

**Figure 2:**
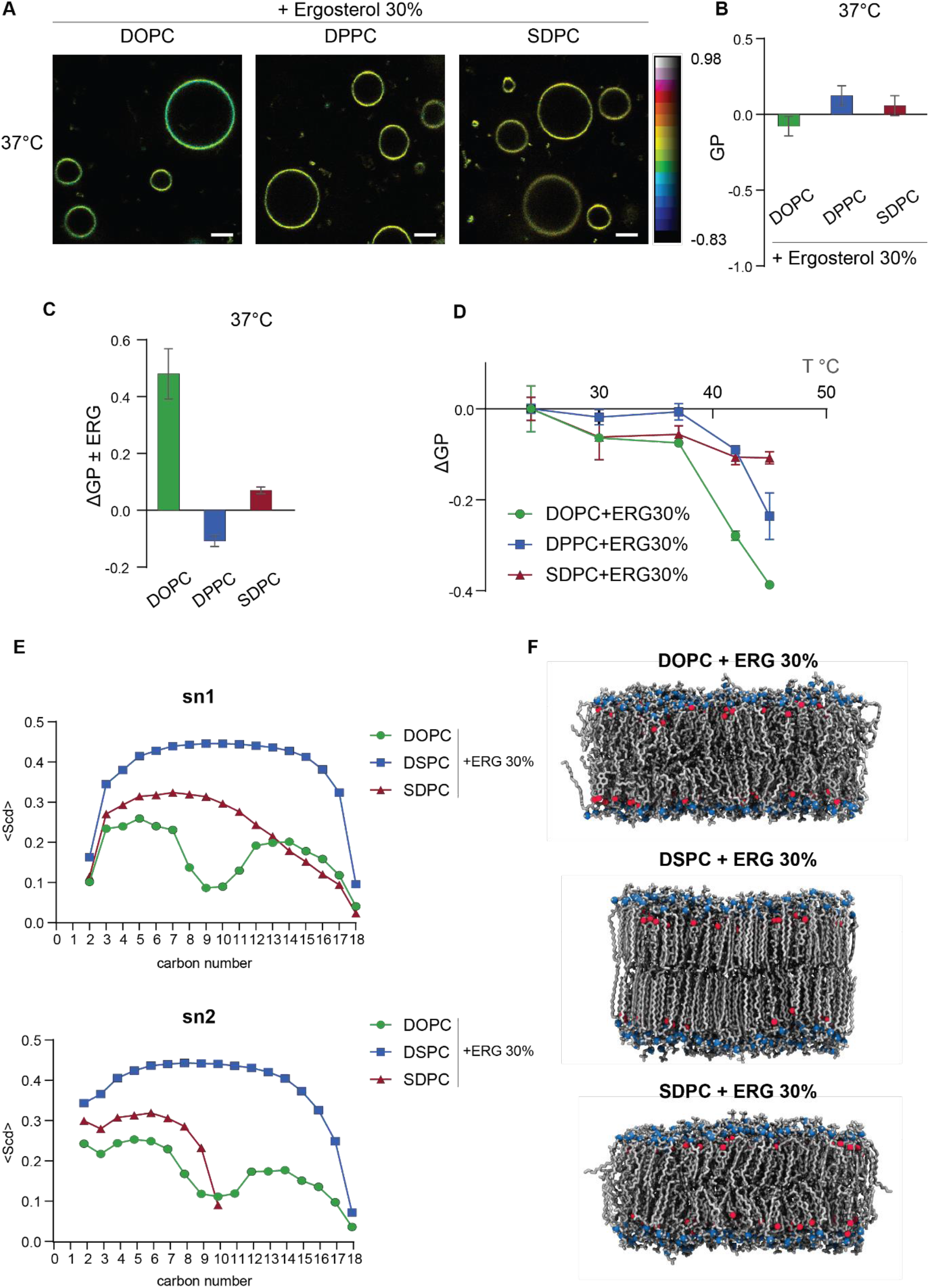
Lipid order of asymmetric lipid bilayers containing 30% ergosterol. A) Representative GP images of asymmetric and symmetric GUVs. Color bar designates GP values. Scale bar, 5 μm. B) Quantification of membrane lipid order in bilayers containing 30% ergosterol. C) Comparison bilayers containing 30% ergosterol relative to pure bilayers showing the minimal effect of ergosterol on SDPC bilayers. D) Variation of GP values with temperature over the range 21°C-45°C showing how SDPC is best able to maintain membrane fluidity. E) Deuterium NMR order parameter for the sn-1 chain (upper graph) and sn-2 (lower graph) for each bilayer. F) Snapshots of atomistic MD simulations of the three bilayers.

### Asymmetric lipids impart bilayers with novel biophysical properties; maintaining high area per lipid and high lipid diffusion despite their lack of acyl chain double bonds

Imaging and simulation data have shown membranes formed from asymmetric phospholipids have intermediate lipid order. Unlike membranes formed from symmetric lipids, they do not require sterol to achieve that membrane order and maintain those properties over a wide range of physiological temperatures. We, therefore, used atomistic MD simulations to test other biophysical membrane properties including lipid packing, membrane thickness and lipid diffusion. Data shows that membranes formed from saturated asymmetric phospholipids have a lipid packing density similar to those formed from unsaturated, symmetric lipids such as DOPC (Figure 3A). Similarly, the data shows that lipid diffusion in SDPC bilayers is significantly higher than that observed in bilayers formed from saturated lipids both in the presence and absence of ergosterol. Diffusion is more akin to that observed for DOPC bilayers, despite the lack of tail unsaturation (Figure 3B).

**Figure 3:**
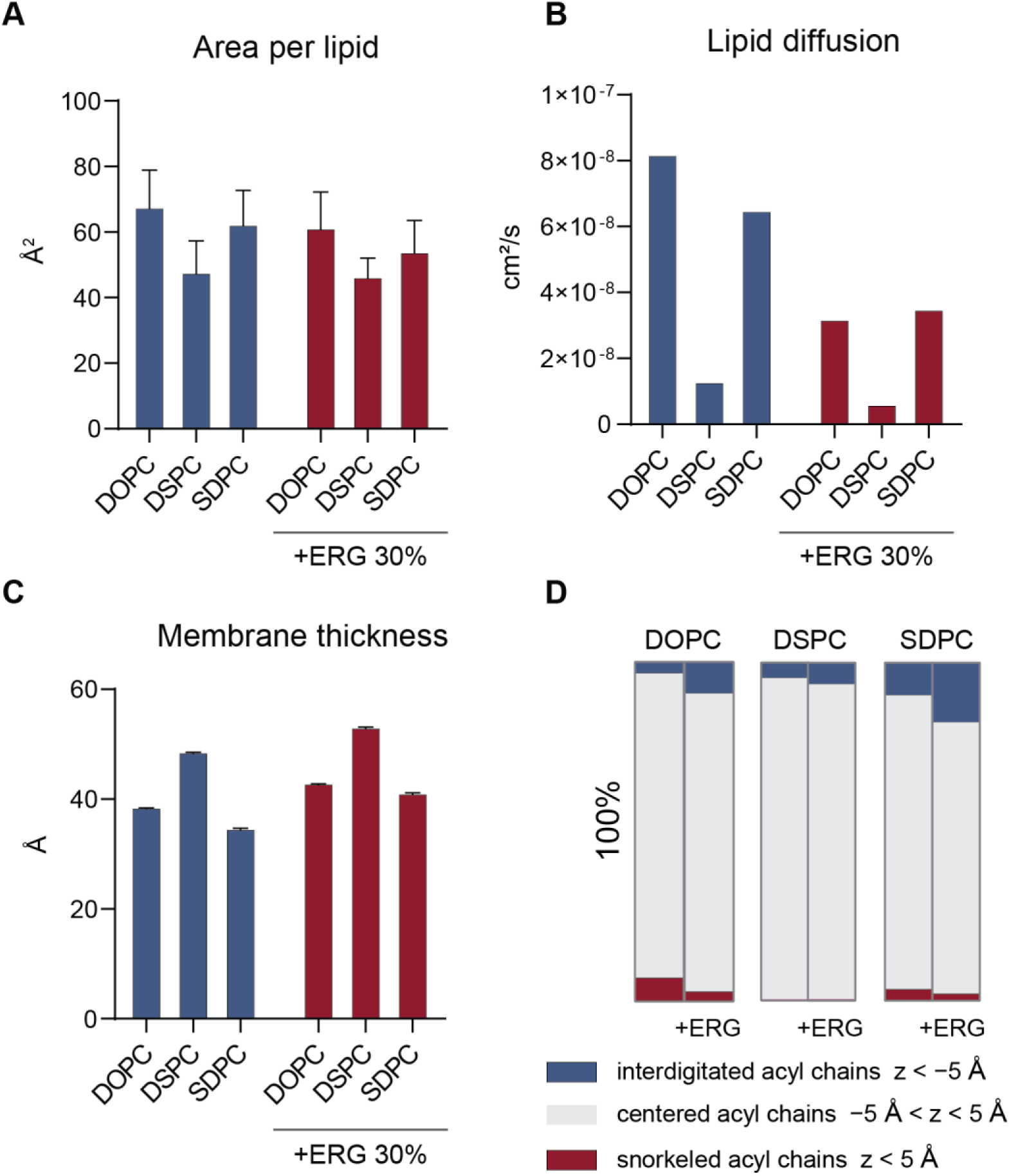
Biophysical properties of bilayers composed of asymmetric phospholipids, extracted from atomistic MD simulations. A) Area per lipid, B) Lipid diffusion coefficient, C) Bilayer thickness and D) Percentage of the terminal carbons in each of the sn acyl chains that interdigitate (z < −5 °A), snorkel (z > 5 °A) and are found in the bilayer center (−5 ° A < z < 5 °A) for each of the simulated bilayers

Finally, asymmetric SDPC bilayers, at only 3.5 nm (4.1 nm in the presence of ergosterol), are significantly thinner than bilayers composed of either saturated (4.9 nm and 5.2 nm) or unsaturated (3.9 nm and 4.0 nm) symmetric lipids, both in the absence and presence of ergosterol (Figure 3C). There are two reasons for this thinning, SDPC bilayers show high levels of snorkeling (3.3% and 1.9% of lipids without/with ergosterol) (Figure 3E), but more significantly, SDPC bilayers show very high levels of interdigitation between the two leaflets (9.7% and 17.75% of lipids). This compared to only 5.0% and 6.8% for saturated symmetric lipids and 3.5% and 9.4% for unsaturated symmetric lipids (Figure 3D).

## Discussion

The fission yeast species, *S. japonicus* produces abundant amounts of asymmetric phospholipids with two saturated acyl chains that differ in length by 8 carbon atoms in place of the more typical symmetric phospholipids present in mammalian cells and even in the closely related sister species and model organism *S. pombe* (Makarova et al., 2020). It is possible that these lipids impart novel biophysical characteristics to bilayers, but these have not previously been studied.

Here, we combined advanced fluorescence microscopy using environmentally sensitive membrane probes and atomistic MD simulations to investigate the novel biophysical properties of bilayers composed of these unusual lipids for the first time. Our data indicate that while membrane lipid order for asymmetric lipid bilayers is intermediate between those composed of either unsaturated or saturated symmetric lipids, they are able to better maintain membrane order over a wider range of physiological temperatures, and crucially above 42°C. It was also notable that the presence of ergosterol – the primary sterol found in fungi, had a smaller effect on asymmetric than symmetric lipids and, importantly, that ergosterol was not required to maintain bilayer fluidity. We therefore conclude that lipid tail asymmetry may be a novel mechanism for modulating membrane fluidity and membrane order in the absence of acyl tail unsaturation and absence of sterols.

*S. japonicus* has previously been shown to survive and proliferate at wider temperature range (up to 42°C), unlike it’s closely related sister species *S. pombe* (Klar, 2013). This data is now consistent with a model by which *S. japonicus* can maintain membrane lipid order above 42°C due to the presence of these asymmetric phospholipids. Interestingly *S. japonicus* grows similarly under anaerobic or aerobic conditions, in contrast to *S. pombe* or other yeast species such as budding yeast *S. cerevisiae,* which exhibit better growth under oxygen supplemented conditions and grow poorly in its absence (Kaino et al., 2018). It is possible that *S. japonicus* evolved a mechanism to adapt to low oxygen consumption and respiration deficiency through a range of adaptive metabolic changes including lipid synthesis toward a high abundance of asymmetric saturated phospholipids. Synthesis of both sterols and unsaturated fatty acids require oxygen, and therefore organisms that can tolerate anaerobic conditions are auxotrophic for sterols and unsaturated fatty acids and many have evolved a mechanism of enhanced uptake of exogenous supply in hypoxic conditions (Reiner et al., 2005). These molecules are key modulators of membrane fluidity and so their absence could partly explain the inability of *S. pombe* and *S. cerevisiae* to grow in these conditions. Interestingly, asymmetric lipids now enter as a novel mechanism to allow *S. japonicus* to survive in such an environmental niche, conferring the ability to modulate and maintain membrane lipid order even in the absence of acyl tail unsaturation or membrane sterols. Consistent with this idea, *S. japonicus* membranes are lower in sterol content compared to *S. pombe* (Makarova et al., 2020), perhaps indicating the necessity of a distinct mechanism to regulate membrane order under a variety of growth conditions and explaining the enhanced survival of *S. japonicus* in both high temperature and hypoxic environments.

MD simulation data shows that the membranes composed of asymmetric lipids are thinner compared to symmetric bilayers. This could also explain the observed significant enrichment of relatively short predicted single transmembrane helices in *S. japonicus* compared to *S. pombe* that doesn’t produce abundant asymmetric lipids (Makarova et al., 2020). Matching of transmembrane domain structure with bilayer properties has recently been shown (Lorent et al., 2020) and may indicate that proteins with shorter transmembrane helices co-evolved with asymmetric lipids (Makarova and Owen, 2020).

The novel biophysical properties imparted to bilayers by unusual asymmetric phospholipids might have a wide range of fascinating consequences and applications. For example, they may represent a more general adaptation to hypoxic environments – in microorganisms, but also mammalian cells such as in solid cancer environments. Industrial use of liposomes (such as for drug delivery) can require specific membrane fluidity which can be difficult to maintain due to oxidation of acyl tail double bonds. We hypothesize that the use of asymmetric lipids for these applications might prove beneficial. Finally, medium chain fatty acids (MCFAs) such as the 10 carbon sn-2 tail of SDPC are an important industrial component of many products from food to cosmetics (Sarria et al., 2017), and are often acquired from environmentally unsustainable sources such as palm oil. We propose *S. japonicus* as a potentially important novel industrial microbe which may prove to be a sustainable source of such molecules.

## Acknowledgments

We thank the fluorescence microscopy imaging facility at the University of Birmingham. Via our membership of the UK’s HEC Materials Chemistry Consortium, which is funded by EPSRC (EP/L000202/1, EP/R029431/1), this work used the ARCHER UK National Supercomputing Service (http://www.archer.ac.uk) and the UK Materials and Molecular Modelling Hub (MMM Hub) for computational resources, which is partially funded by EPSRC (EP/P020194/1) to carry out the MD simulations reported in this manuscript. P.S. acknowledges the funding provided by the EPSRC DTP Studentship Block Grant. D.M.O. acknowledges funding from BBSRC grant BB/R007365/1.

## Author contributions

P.S. and C.D.L. performed MD simulations and analysis. M.M. and D.M.O performed microscopy experiments and analysis. P.S., C.D.L., M.M. and D.M.O. wrote the manuscript. M.M. conceived the work.

